# Impacts of age on the gut microbiota in captive giant pandas

**DOI:** 10.1101/2023.02.02.526921

**Authors:** Huixin Li, Kangning Lu, Guo Li, Ti Li, Le Zhang, Chao Li, Qingyang Xie, Huaiting Liu, Xinxing Zhang, Minghao Gong, Gang Liu, Guiquan Zhang

## Abstract

The gut microbiota is the most complex and most abundant symbiotic microbial ecosystem in animals. Aging is one of the main factors that cause gut microbiota structure changes, and the relationship between age and the gut microbiota in the giant panda has been a key focus of attention. The giant panda has a specialized diet of bamboo, and it relies on the microbiota that colonizes its gut to complete digestion. However, there is no in-depth understanding of the changes in the gut microbiota across the lifespan of giant pandas. Here, we identified the differences in the gut microbiota between four age groups (cubs, juveniles, adults, and geriatrics) using 16S rRNA gene high-throughput sequencing on an Illumina MiSeq platform. The results revealed that Firmicutes (mean±SD: 65.45±30.21%; range: 0.91–99.62%) and Proteobacteria (mean±SD: 31.49±27.99%; range: 0.26–85.35%) were the dominant phyla. The relative abundance of *Escherichia-Shigella* was high in both the cubs and juveniles. It is interesting to note that the adults had the highest richness and lowest diversity, while the cubs had the opposite. In summary, our study indicates that the gut microbial community composition, abundance, and functional pathways differ across four age groups of giant pandas. Exploring the influence of age, an endogenous influencing factor, on gut microbes provides basic scientific data for monitoring gut microbial dynamics and formulating gut microbial health management approaches, thereby improving the protection of giant pandas.

## 1 Introduction

In recent years, the symbiotic relationships between hosts and their gut microbiota have become a research hotspot. The gut microbiota plays vital roles in food digestion, immune response, allergic response, nutrient absorption, and energy metabolism (Ramachandra et al., 2018, Maifeld et al., Rowland et al., 2018, Näpflin and Schmid-Hempel, 2018). The colonization and diversity of the gut microbes are affected by both internal and external factors (Thakur et al., 2014). Internal factors include age, sex, and host genetic background, while external factors include geographic location, environmental conditions such as climate, diet, deworming history, and drug use (Amato et al., 2013, Candela et al., 2014, Carmody et al., 2015, Zhang et al., 2021, Linnenbrink et al., 2013, Holmes et al., 2012, Spor et al., 2011). For endangered species, gut microbes directly affect the long-term development of the population. Thus, much attention has been focused on the gut microbiota of endangered species and the influencing factors. The factors that influence the gut microbiota of endangered species such as Amur tiger, forest musk deer (*Moschus berezovskii*), and great apes (Ning et al., 2020, Hu et al., 2018, Hicks et al., 2018) have been extensively studied.

The giant panda (*Ailuropoda melanoleucthe*) is a key icon in the field of conservation biology and is well known for its specialized and exclusive diet of bamboo (Wei et al., 2012, Wei et al., 2015). The mechanism underlying how the giant panda is adapted to a bamboo diet is a hot topic. The 16S rRNA diversity assessment method and metagenomics have been used to analyze the role of the giant panda’s gut microbiota in the digestion and hydrolysis of bamboo. Research has revealed that the gut microbiota possesses cellulose-degrading genes, which play an essential role in digesting cellulose and hemicellulose (Hirayama et al., 1989, Li et al., 2010, Zhu et al., 2011, Wei et al., 2015). To reveal the main drivers of the gut microbiota composition in giant pandas and other specialized species, the phylogeny and diet of giant pandas have been studied. The giant panda has a carnivore-like gut microbiota that is less biodiverse than that of herbivores and does not correspond to the composition of the red panda’s gut microbiota (Li et al., 2015, Xue et al., 2015). Furthermore, research has shown that several endogenous and environmental factors can impact both the composition and functionality of the gut microbiota in giant pandas (Williams et al., 2013, Zhao et al., 2017, Wu et al., 2017, Scoma et al., 2020).

However, only a few studies have investigated the effects of age on the gut microbiota in giant pandas. Studies report different or even inconsistent results, and no study has covered all age groups across the lifespan of giant pandas. To reveal whether age is the main driver of changes in the gut microbiota of giant pandas, it is necessary to analyze the changes across the lifespan. As gastrointestinal tract disease threatens the health of giant panda populations in ex-situ conservation programs, it is key to explore the influence of age on gut microbes. This will provide basic scientific data for monitoring gut microbial dynamics and formulating gut microbial health management approaches, thereby improving the protection of giant pandas.

In 2021, the number of captive giant pandas in China reached 673. There has been a stable increase in population size over the last several decades, owing to scientific and effective management of breeding and nursing. Different management approaches are used for different age groups of giant pandas, with age-specific management being based on scientific knowledge. The gut microbiota is an important indicator for use when assessing feeding and breeding, and even when evaluating giant panda reintroduction projects (Tang et al., 2020), so it is necessary to explore how the gut microbiota differs among age groups. In this study, fecal samples were collected from captive pandas, which were divided into four age groups: cubs (0–1.5 years old), juveniles (1.5–5.5 years old), adults (5.5–20 years old), and geriatrics (>20 years old) (Jinchu, 2011). The 16S rDNA V3–V4 region underwent high-throughput sequencing, and the gut microbiota composition, diversity, and function of giant panda cubs, juveniles, adults, and geriatrics were compared. According to the characteristics of the gut microbiota at different stages, different strategies are suggested to facilitate the management of different age groups in the captive giant panda population.

## 2 Material and Methods

### 2.1 Sample collection

A total of 56 fresh fecal samples were collected in Sichuan Province, China, in April 2017. The feces (N=56) were collected from the China Conservation and Research Center for Giant Panda Bifengxia Base (n=24), Dujiangyan Base (n=22), Shenshuping Base (n=9), and Chongqing Zoo (n=1). For pandas of the same age, the four breeding centers have very similar management processes and standard food composition, and all the pandas have access to clean water. Besides, the four breeding centers have very similar in geographical distribution and climate. Therefore, it is believed that different breeding centers do not have significant effects on gut microbes。 Among the pandas we collected feces from, there were 18 cubs (aged <9 months; not weaned; 7 females, 4 males, and 7 unreported), 12 juveniles (aged 1.5–4 years; 6 females and 6 males), 12 adults (aged 5–19 years; 6 females and 6 males), and 14 geriatrics (aged >20 years; 3 males and 11 females) (Table S1). The cubs’ diet consisted mainly of dairy products and young bamboo leaves. During the cub stage, some cubs ate dairy products, while others also ate bamboo. The diets of the juveniles, adults, and geriatrics were basically the same, mainly consisting of bamboo leaves, bamboo stems, bamboo shoots, and panda cake (mix the fruits and vegetables). Samples of fresh feces were collected immediately after natural defecation and placed in sterile plastic containers. They were initially stored in dry ice at the breeding centers and then transport to the lab within 24h. Samples of fresh feces were stored at −80°C in the lab less than 24h.

All sample collection procedures followed the guidance of the Sichuan Animal Society Ethics Committee and had no impact on animal health.

### 2.3 DNA extraction

The environmental DNA (eDNA) in the feces was extracted using an E.Z.N.A.^™^ Mag-Bind Soil DNA Kit (OMEGA, USA) according to the manufacturer’s instructions.

The extracted genomic DNA was assessed using 1% agarose gel electrophoresis to check the integrity and concentration. The genomic DNA was then stored at −80°C.

### 2.4 PCR amplification and sequencing

A Qubit 2.0 DNA Detection Kit (Life Technologies, Carlsbad, USA) was used to accurately quantify the genomic DNA to determine the amount of DNA to be used for PCR. The V3–V4 region of 16S rRNA was amplified by PCR. The PCR primers involved a V3–V4 universal primer (341F and 805R) fused with a unique barcode primer: forward primer: CCTACGGGNGGCWGCAG (341F); reverse primer: GACTACHVGGGTATCTAATCC (805R).

To improve the primer efficiency, two rounds of PCR amplification were carried out. The first PCR amplification system volume was 30μL, comprising 15μL 2×Taq master Mix, upstream and downstream primers (1μL each, 10μM), 10-20ng DNA template, add the H2O to 30μL. The PCR amplification conditions were as follows: 94°C for 3min; 5 cycles of 94°C for 30s, 45°C for 20s, and 65°C for 30s; 20 cycles of 94°C for 20s, 55°C for 20s, and 72°C for 30s; followed by 72°C for 10min. The second PCR amplification system was the same to the first. The PCR amplification conditions were as follows: 95°C for 3min; 5 cycles of 94°C for 20s, 55°C for 20s, and 72°C for 30s; followed by 72°C for 5min.

The PCR products were subjected to agarose gel electrophoresis. They were purified using 0.6×Agencourt AMPure XP (Beckman Coulter, Brea, CA) according to the manufacturer’s instructions. They were then sequenced by Shanghai Biotechnology Corporation on a MiSeq PE300 sequencing system (Illumina, San Diego, USA).

### 2.5 Statistical and bioinformatics analyses

#### 2.5.1 Analysis of gut microbiota composition in giant pandas

The MiSeq sequences were optimized by quality control, splicing, and chimera removal. FLASH (version 1.2.11, https://ccb.jhu.edu/software/FLA) and FASTP (version 0.19.6, https://github.com/OpenGene/fast) were used to filter, splice, remove the barcode and primers at both ends, and finally obtain the representative sequences. The sequences were divided into operational taxonomic units (OTUs), based on a 97% similarity threshold, using UPARSE (version 7.0.1090, http://drive5.com/uparse/). To classify the representative OTU sequences, the RDP Classifier (version 2.11, http://sourceforge.net/projects/rdp-classifier/) algorithm (default confidence threshold, 0.7) was used, using Silva (Release138, http://www.arb-silva.de) for species annotation. Mothur (version 1.30.1, http://www.mothur.Org/wiki/Schloss_SOP#Alpha_diversity) was used for the alpha diversity index analysis of all samples. Observed species index, Chao1 index, Shannon index, Simpson index, abundance-based coverage estimator (ACE) index, and Good’s coverage index were used to evaluate alpha diversity.

#### 2.5.2 Age-related differences in gut microbiota

For beta diversity, non-metric multidimensional scaling (NMDS) analysis involving the OTU-based unweighted UniFrac distance matrix using QIIME (version. 1.1.9.1, http://qiime.org/install/index.html) was conducted. One-way analysis of similarity (ANOSIM) was used to test whether the difference between groups was significantly greater than the difference within groups, to assess whether the grouping was meaningful. Nonparametric permutational multivariate analysis of variance (PERMANOVA) based on the Bray–Curtis matrix was used to analyze whether groups of samples were significantly different based on age. LEfSe software (http://huttenhower.sph.harvard.edu/galaxy/root?tool_id=lefse_upload) was used to conduct a linear discriminant analysis (LDA) effect size (LEfSe) analysis and LDA. LEfSe was used to identify the gut bacteria (based on OTUs) that significantly characterize the four groups. LDA was used to reduce the dimensionality of the data to assess the influence of significantly different species. Adonis, a nonparametric implementation of a permutational analysis of variance, was used to estimate the statistical significance between groups. The distance between two samples was first calculated using the distance algorithm (default Bray-Curtis), and then all distances were sorted from smallest to largest and the p-value is calculated. PLS-DA, a variant of the partial least squares (PLS) regression, used a regression method for classification.

#### 2.5.3 Prediction of gut microbial functions in the different age groups

PICRUSt is a bioinformatics tool that utilizes marker genes to predict gene family abundance in environmental DNA samples for which only marker gene data are available. The OTU abundance table was normalized using PICRUSt (version.1.1.0, http://picrust.github.io/picrust/). The OTUs were annotated with Clusters of Orthologous Groups (COG) family information and KEGG Ortholog (KO) information using the evolutionary genealogy of genes: Non-supervised Orthologous Groups (EggNOG; http://eggnog.embl.de/) database and Kyoto Encyclopedia of Genes and Genomes (KEGG) database (http://www.genome.jp/kegg/). The abundance of each COG and KO was then calculated.

#### 2.5.4 Statistical analysis

SPSS (version 20) was employed to test the significance of differences in the relative abundances of gut bacteria and alpha diversity indexes. The significance of differences between multiple groups was tested by the Kruskal–Wallis H test, while the significance of differences between two groups was tested by the Mann–Whitney U test. Other statistical calculations were conducted using the statistical programming language R (version 3.3.1). Charts were plotted using the R packages ‘vegan’ and ‘ggplot2’.

##### Data availability

Shotgun metagenomic reads generated for this study were uploaded to NGDC under the GSA no. CRA009434 (https://ngdc.cncb.ac.cn/search/?dbId=&q=CRA009434).

## 3 Results

### 3.1 Gut microbiota in different age groups

Among the fecal samples of 56 pandas, there were 4,066,208 optimized 16S rRNA gene sequences, involving 1,736,583,084 optimized bp. A mean±SD of 72610.86±11,679.57 sequences (range: 42,970–93,193) was obtained per sample.

The Good’s estimates for the 56 samples were >98% (Figure S1A), suggesting that >98% of the diversity estimated in the samples was recovered. The results of the species-accumulation curves indicate that the number of OTUs leveled off as the sample size increased and the sample size was adequate (Figure S1B). There were 1,140 OTUs (97% similarity threshold) with the minimum number of sample sequences. The mean number of OTUs per sample was 105 (range: 35–759). A total of 375 OTUs were found in the cub group, 304 in the juvenile group, 347 in the adult group, and 343 in the geriatric group. However, there were 759 OTUs in sample C4 in the cub group, which was much higher than in the other 55 samples. Therefore, in the subsequent analysis, C4 was removed. Rarefaction curves of Shannon index values indicated that the bacterial diversity of each sample was fully measured at the sequencing depth used (Figure S1C). Rank abundance curves (indicating species richness and evenness), which tend to be horizontal if there is high evenness, indicated that the sequencing depth was sufficient to represent the gut microbiota diversity in the four age groups (Figure S1D).

The OTUs were divided into 27 phyla and 374 genera. At the phylum level (Table S2), Firmicutes and Proteobacteria were dominant, accounting for 65.45±30.21% (0.91–99.62%) and 31.49±27.99% (0.26–85.35%) of the total sequences, respectively. The subdominant phyla were Bacteroidetes (1.59±6.91%; 0–46.43%), Cyanobacteria (0.79±1.84%; 0–12.8%), and Actinobacteria (0.63±1.71%; 0–8.49%). At the genus level (Table S3), the top two were *Streptococcus* (47.43±40.03%; 0.02%–47.43%) and *Escherichia-Shigella* (27.22±26.86%; 0.03%–27.22%). Other genera with a relative abundance >1% were *Lactobacillus* (7.98±16.73%; 0–7.98%), *Clostridium_sensu_stricto_1* (4.2±7.88%; 0–4.2%), *Pseudomonas* (1.49±8.04%; 0–1.49%), *Turicibacter* (1.27±3.39%; 0–1.27%), and *Megasphaera* (1.26±3.8%; 0–1.26%).

There were differences in gut microbiota composition among the four age groups (Table S4). The cubs had the most OTUs (383) and the juveniles had the fewest OTUs (337) (Figures 1A and 1B). The cubs’ and juveniles’ dominant phyla were Proteobacteria (cubs: 51.83±22.6%; juveniles: 51.83±22.6%) and Firmicutes (cubs: 42.51±24.82%; juveniles: 42.51±24.82%). The cubs’ dominant genera were *Escherichia-Shigella* (46.8±25.4%) and *Lactobacillus* (24.8±22.4%). The juveniles’ dominant genera were *Streptococcus* (63.2±30%) and *Escherichia-Shigella* (18.8±18%). The adults had 381 OTUs; the dominant phyla were Firmicutes (85.32±22.26%) and Proteobacteria (12.6±21.78%) and the dominant genera were *Streptococcus* (79.1±25.4%) and *Escherichia-Shigella* (11.5±21.5%). The geriatrics had 386 OTUs; the dominant phyla were Firmicutes (71.53±29.11%) and Proteobacteria (27.37±29.07%) and the dominant genera were *Streptococcus* (62.7±34.1%) and *Escherichia-Shigella* (24.2±27.1%).

**Figure 1.**
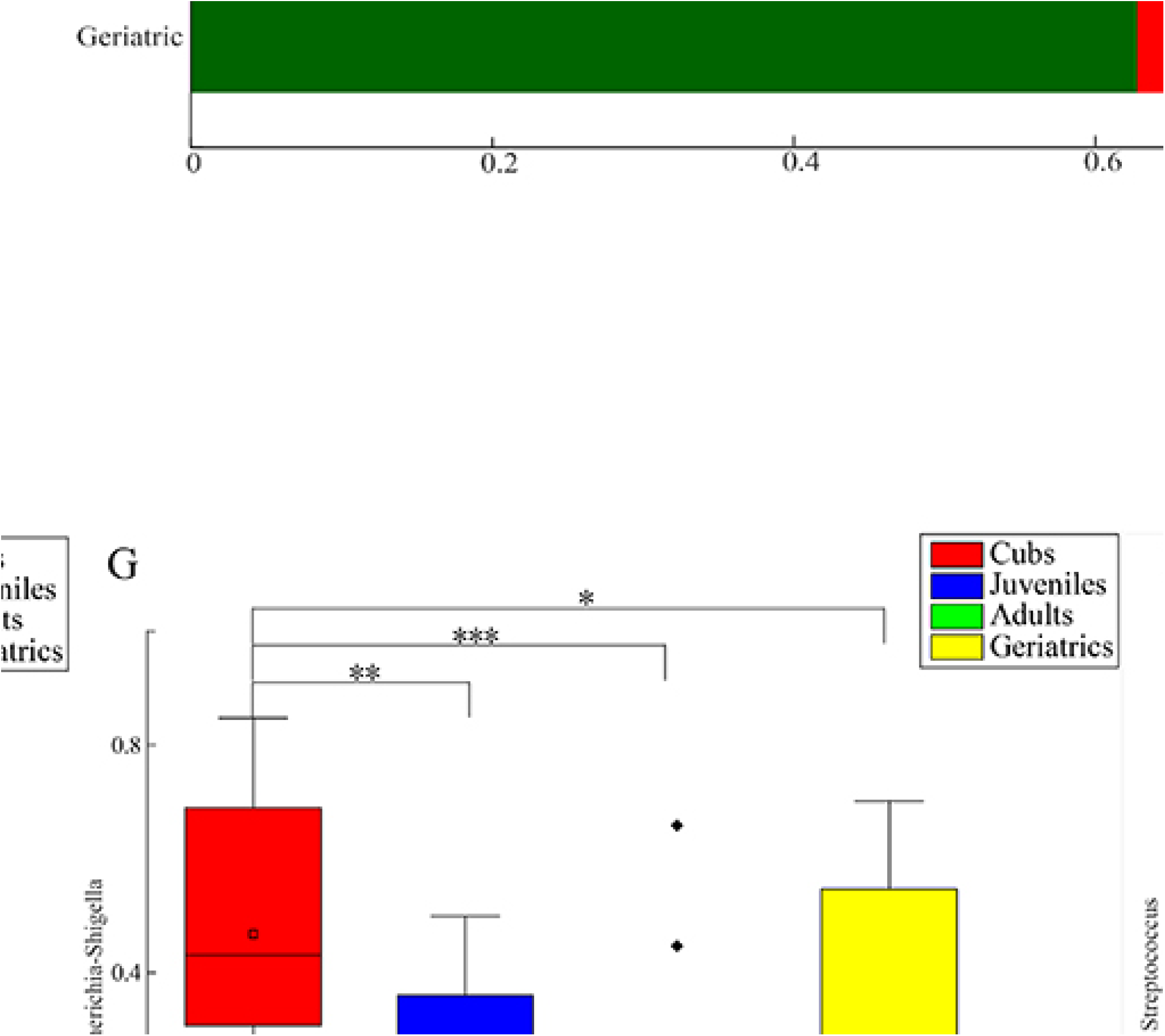
(A) and (B) Histograms of relative abundances of gut bacteria in four age groups of giant pandas. (C–I) Box plots of relative abundances of gut bacteria with significant differences among four age groups of giant pandas at phylum and genus levels. (J–N) Box plots of alpha diversity of gut microbiota among four age groups of giant pandas.

At the phylum level(Table S5), the differences in the relative abundance of Firmicutes, Cyanobacteria, Proteobacteria, and Actinobacteria among the four groups were extremely significant (P≤0.001) (Figures 1C–E). The relative abundances of Firmicutes and Cyanobacteria were highest in the adults, while the relative abundances of Proteobacteria and Actinobacteria were highest in the cubs. At the genus level(Table S6), the relative abundance of *Streptococcus* was highest in the adults, while *Escherichia-Shigella* and *Lactobacillus* were highest in the cubs (Figures 1H–I).

The alpha diversity indexes differed among the age groups(Table S7). The Shannon index was significantly higher in the cubs (indicating low diversity) than the others (P≤0.01), while the adults had the lowest value (Figures 1J–N). The Simpson index was significantly higher in the adults (indicating high diversity) than the others (P≤0.01), while the cubs had the lowest value. The ACE index and Chao 1 index were also significantly different among the age groups (P≤0.01), with the adults having the highest and the cubs having the lowest values.

### 3.2 Core gut microbiota composition in different age groups

The graph showing the pan analysis curves (Figure S5) exhibited an upward trend for each age group, but the graph showing the core analysis curves exhibited a horizontal line for each age group (between 10 to 20). This indicates that increasing the sample size may increase the total number of OTUs, but not the number of core OTUs. The number of OTUs shared by all pandas in each group was 124 (Figure 2A). These core OTUs which are shared by all giant pandas belonged to 4 phyla and 7 genera, mainly including Firmicutes (61.96%), Proteobacteria (35.99%), *Streptococcus* (47.58%), and *Escherichia-Shigella* (31.48%) (Figures 2B and 2C).

**Figure 2.**
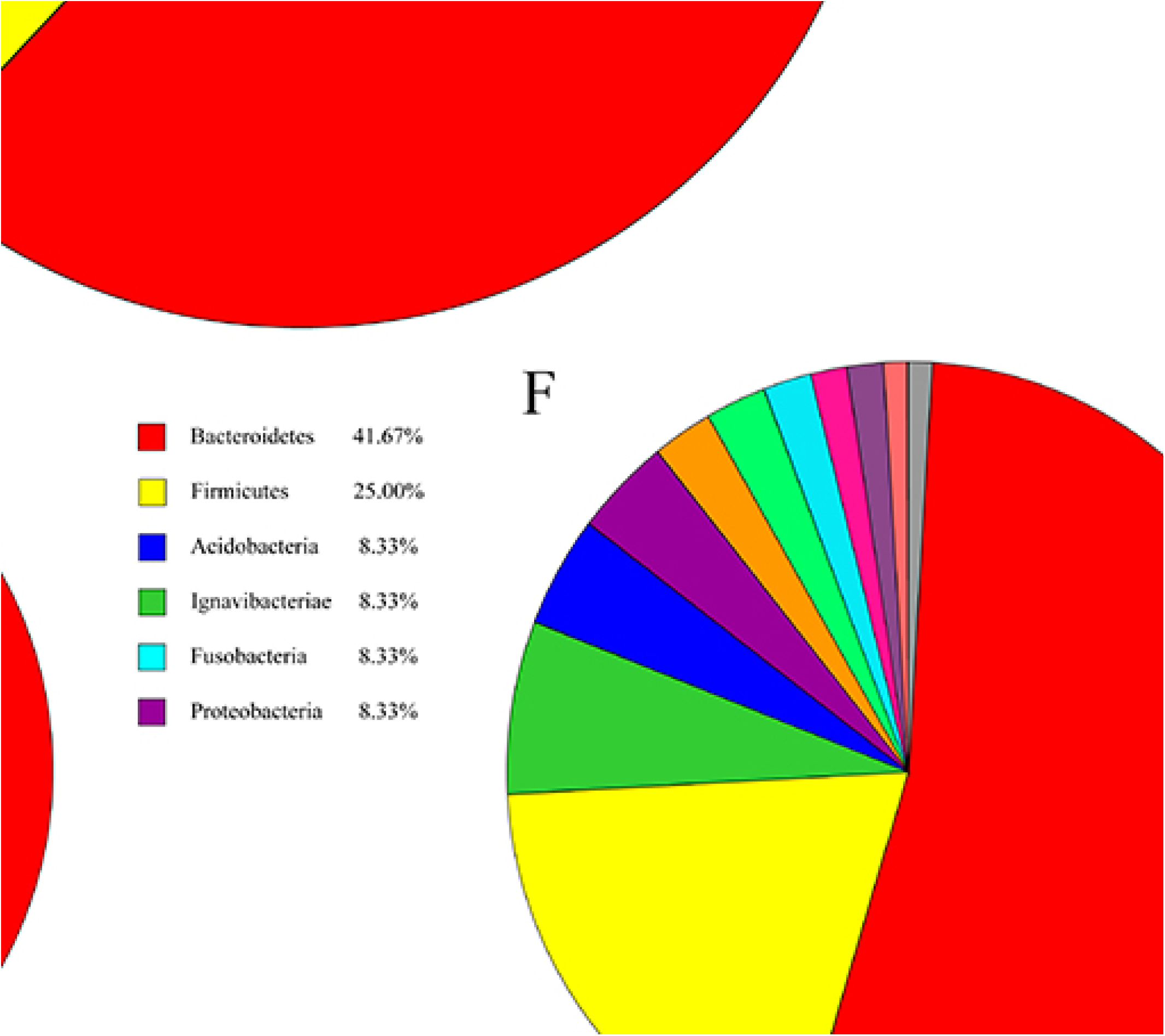
(A) UpSet plot of the distribution of shared and unique OTUs (97%similarity threshold) among four age groups of giant pandas. Pie charts of the core bacteria among four age groups of giant pandas at the (B) phylum and (C) genus levels. Pie charts of the unique bacteria at the (D–G) phylum and (H–K) genus levels in the cubs, juveniles, adults, and geriatrics, respectively.

The numbers of OTUs (relative abundance ≥1%) unique to all pandas in the cub, juvenile, adult, and geriatric groups were 117, 44, 52, and 61, respectively. In the adults, the unique OTUs belonged to 10 phyla and 19 genera, mainly including Firmicutes (53.66%), Bacteroidetes (19.5%), *Ezakiella* (15.61%), and *Dialister* (13.66%) (Figures 2F–G). In the juveniles, the unique OTUs belonged to 6 phyla and 11 genera, mainly including Bacteroidetes (41.67%), Firmicutes (25%), Parabacteroides (16.67%), and *norank_f_Saprospiraceae* (8.33%) (Figures 2E and 2I). In the geriatrics, the unique OTUs belonged to 11 phyla and 7 genera, mainly including Parabacteroides (28.57%), Firmicutes (21.43%), *Alteromonas* (14.29%), and *Shuttleworthia* (14.29%) (Figures 2G and 2K). In the cubs, the unique OTUs belonged to 12 phyla and 13 genera, mainly including Firmicutes (66.18%), Actinobacteria (25%), Bacteroidetes (8.82%), *Megasphaera* (39.71%), and *Lactobacillus* (16.18%) (Figures 2D and 2H).

### 3.3 Age-related differences in gut microbiota structure

The heatmaps of the gut microbiota in the four age groups indicated that the cubs significantly differed from other groups. The juveniles and geriatrics were clustered together. The significant differences in the relative abundances of the gut microbiota phylum- and genus-level communities were consistent with the overall findings shown in the heatmaps (Figures 3A and 3B).

**Figure 3.**
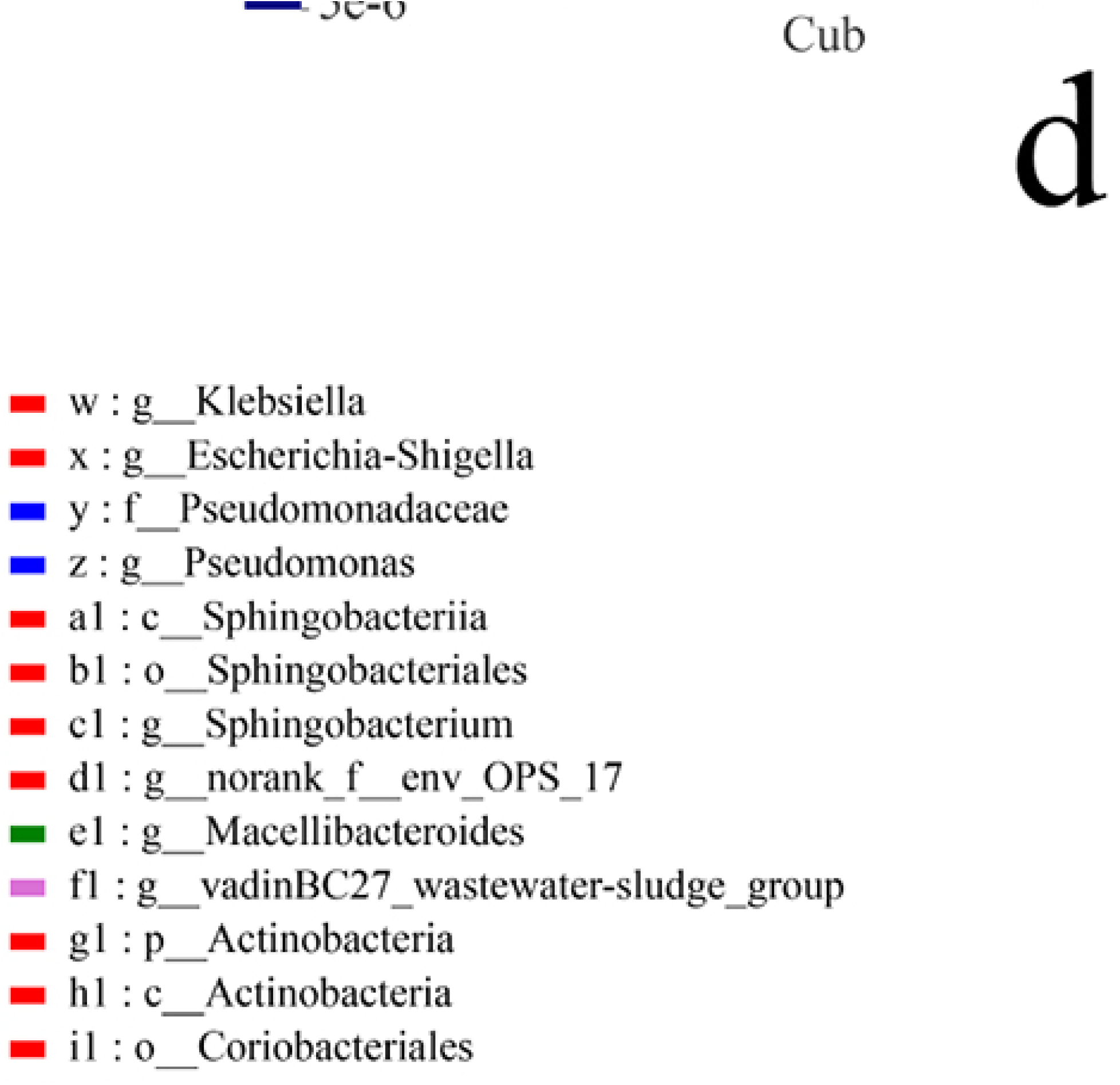
(A) and (B) Heatmaps of giant panda gut microbiota community composition at phylum and genus levels. (C) Differences in taxa abundances among four age groups identified by LEfSe. Colored circles and shaded areas are used to highlight biomarker taxa (cubs: red; juveniles: blue; adults: green; geriatrics: purple). The diameter of each circle reflects the relative abundance of taxa in the community. (D) Differences in taxa abundances among four age groups identified by LDA (cubs: red; juveniles: blue; adults: green; geriatrics: purple), with cutoff value ⩾4.0. (E) Non-metric multidimensional scaling (NMDS) plot of the four age groups based on UniFrac distance. (F) Partial least squares-discriminant analysis (PLS-DA) plot of the four age groups, which explains the different age groups’ gut microbiota. Comp1 and comp2 represent the influencing factors.

Using LEfSe analysis, we were able to identify the gut bacteria that significantly (LDA>4.0 and p≤0.05) characterized the four groups (Figures 3C and 3D). Based on the LDA scores and cladogram assay, the gut bacteria that significantly characterized the cubs were Proteobacteria at the phylum level, Gammaproteobacteria, Negativicutes, and Sphingobacteriia at the class level, Enterobacteriales, Selenomonadales, and Sphingobacteriales at the order level, Enterobacteriaceae, Veillonellaceae, and Lactobacillaceae at the family level, and *Escherichia-Shigella*, *Megasphaera, Sarcina*, and *Lactobacillus* at the genus level (LDA>4.0 and p≤0.05). The corresponding taxa for juveniles were Pseudomonadaceae (family), and *Pseudomonas* (genus) (LDA>4.0 and p≤0.05). The corresponding taxa for adults were Firmicutes (phylum), Bacilli (class), Lactobacillales (order), Streptococcaceae (family), and *Streptococcus* (genus) (LDA>4.0 and p≤0.05). The corresponding taxon for geriatrics was *vadinBC27_wastewater_sludge_group* (genus belonging to the phylum Bacteroidetes) (LDA>4.0 and p≤0.05).

The NMDS analysis indicated that the juveniles, adults, and geriatrics partially overlapped, while the cubs were completely separated from the other groups (R=0.4504, P=0.001) (Figure 3E). ANOSIM showed that the inter-group differences were greater than the intra-group differences (P=0.4504, R=0.001). Adonis (PERMANOVA) showed that the different grouping factors could explain the differences among samples to a high degree and with reliability (R^2^=0.05, P=0.04). Partial least squares discriminant analysis (PLS-DA) showed that comp1 and comp2 could explain the results with a weight ratio of 21.5%. The cubs and adults were separated into two subgroups by comp1, while the juveniles and the other three groups were distinguished by comp2 (Figure 3F). PERMANOVA also explained the relationship between the age groups (R=0.43398, P=0.001).

### 3.4 Prediction of gut microbial functions in the different age groups

The gut microbiota played important roles in amino acid transport and metabolism, carbohydrate transport and metabolism, translation, ribosomal structure, and biogenesis (Figure 4A). One-way ANOVA analysis indicated that the results of KEGG functional enrichment analysis of gut microbes in giant pandas differed among age groups, especially for Human diseases, Cellular processes, Genetic information processing, and Environmental information processing was highly significant (P⩽0.001)(Figure 4B). The heatmaps of the Enzyme in the four age groups indicated that the cubs significantly differed from other groups (Figure 4C). To assess gut microbial functions, all quantified microbial proteins were annotated using the COG database. Among the 10 functions with the highest relative abundance, the differences were highly significant (P⩽0.001) for Function unknown, Carbohydrate transport and metabolism, Translation, ribosomal structure and biogenesis, Inorganic ion transport and metabolism, Cell wall/membrane/envelope biogenesis, Energy production and conversion, Nucleotide transport and metabolism in four age groups, and significant (P⩽0.01) for Carbohydrate transport and metabolism (Figure 4D). Functional analysis of four groups of giant panda gut microbiome was performed with the MetaCyc metabolic pathway database, and the results showed that among the top ten metabolic pathways in terms of enrichment abundance, pyruvate fermentation to isobutanol (engineered), sucrose degradation III (sucrose invertase), peptidoglycan maturation (meso-diaminopimelate containing) and pentose phosphate pathway (non-oxidative branch) were highly significant (P⩽0.001), while CDP-diacylglycerol biosynthesis I, CDP-diacylglycerol biosynthesis II, superpathway of pyrimidine nucleobases salvage, adenosine deoxyribonucleotides de novo biosynthesis II, guanosine deoxyribonucleotides de novo biosynthesis II, and acetylene degradation were significant (0.01≤P<0.001) (Figure 4E).

**Figure 4.**
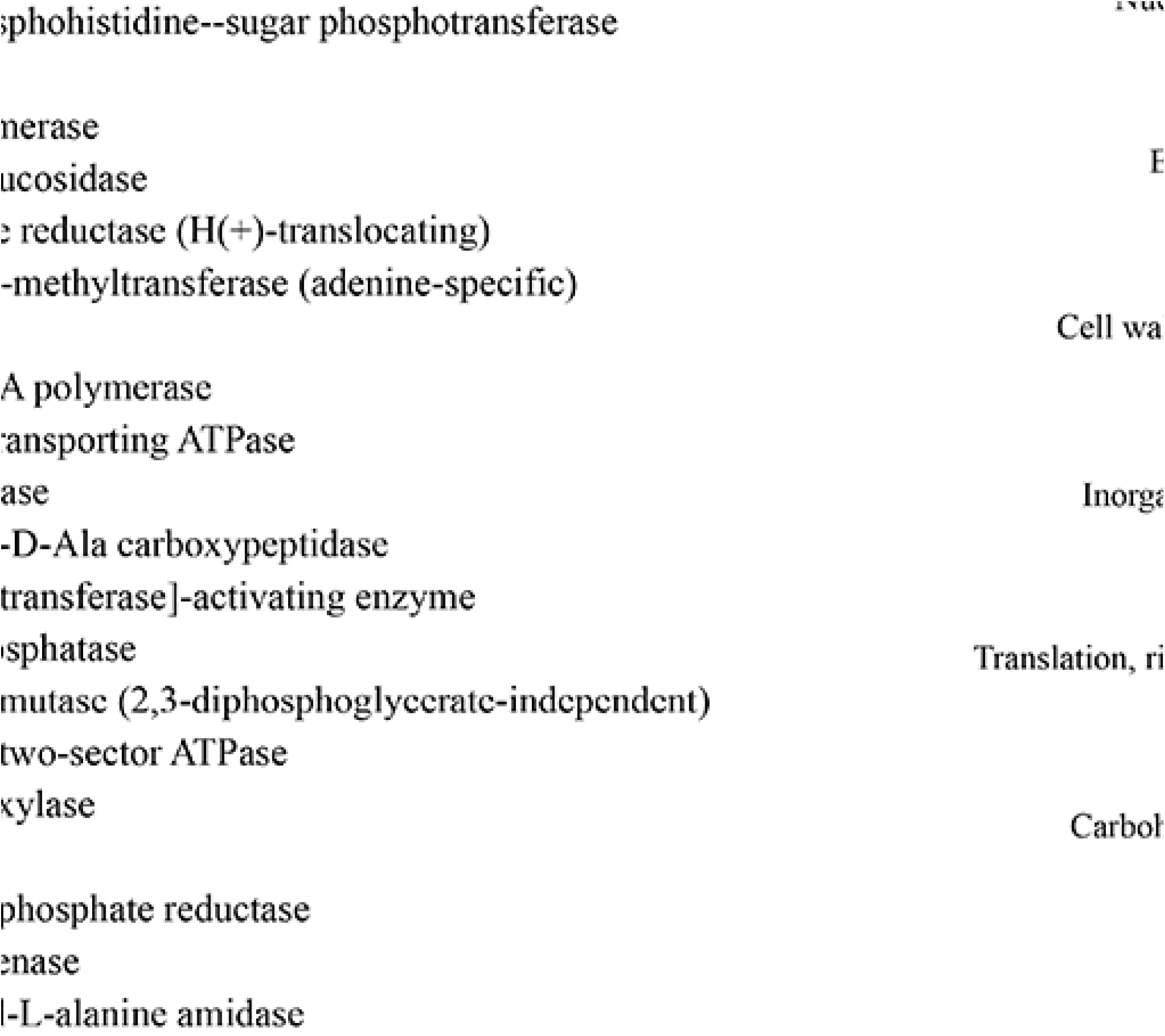
(A) Histogram of Clusters of Orthologous Groups (COG) function classification for each age group, predicted by PICRUSt. (B) One-way ANOVA analysis KEGG functional enrichment of gut microbes in giant pandas differed among age groups. (C) The heatmaps of the Enzyme in the four age groups of giant pandas. (D) One-way ANOVA analysis the top 10 of COG enrichment functions of gut microbes in giant pandas differed among age groups. (E) One-way ANOVA analysis the top 10 of MetaCyc enrichment functions of gut microbes in giant pandas differed among age groups.

## 4 Discussion

In humans, primates, and captive animals, research has shown that age affects gut microbial composition and diversity (Lim et al., 2019, Janiak et al., 2021, Yao et al., 2021). There have also been several studies on the effect of age on the gut microbes of giant pandas. However, these studies on the age-related changes in the gut microbiota of giant pandas have two major limitations: (1) there are no studies on the complete lifespan of giant pandas, as the study subjects usually only belonged to 1–3 age groups, such as newborns and 1-year-olds; juveniles, adults, and geriatrics; and adults and geriatrics (Zhan et al., 2019, Tun et al., 2014) and (2) the sample sizes in the age groups were small (Tun et al., 2014). To determine the relationship between age and the gut microbiota across the giant pandas’ lifespan, we divided the giant pandas into four groups: cubs, juveniles, adults, and geriatrics. This grouping strategy allowed the relationships among gut microbial composition, diversity, and host developmental stages to be effectively evaluated, providing a scientific basis for accurate health prediction and management.

In this study, Firmicutes and Proteobacteria were the dominant gut microbiota. This finding is consistent with previous research on giant panda gut microbial composition (Yan et al., 2021). These bacteria play a crucial role in the digestion of cellulose and hemicellulose in giant pandas (Wei et al., 2015, Zhang et al., 2018b, Fang et al., 2012). However, there are differences in the relative abundances of dominant bacterial groups in different studies (Hirayama et al., 1989). This may be caused by differences in factors such as the captive environment, sampling season, edible bamboo species, and climate (Zhu et al., 2011, Zhang et al., 2018b, Xue et al., 2015, Yan et al., 2021). The dominant species, composition, and structure of the gut microbial community were significantly different between the cubs and the other three groups. At the phylum level, the relative abundances of dominant and subdominant bacteria in the four age groups were significantly different. The relative abundance of Firmicutes and Cyanobacteria were highest in the adults, while the relative abundance of Proteobacteria and Actinobacteria were highest in the cubs. Firmicutes plays a major role in fiber digestion in the guts of humans and other mammals. Our research indicates that as the age of giant pandas increased, Firmicutes in the gut microbiota increased significantly, peaking in adulthood. Although the relative abundance of Bacteroidetes was not significantly different among the four age groups, it non-significantly decreased with age. This indicates that increased age leads to transformation of the gut microbiota composition so that the giant panda is adapted to high-fiber food and hydrolysis can occur efficiently to obtain nutrients to sustain life.

At the genus level, the genera with a relative abundance >1% differed among the four age groups. *Escherichia-Shigella* and *Lactobacillus* made up the majority of gut microbes in the cubs, and other studies have found similar results (Guo M et al., 2019). During early gastrointestinal tract colonization in the cub, the dominance of the cub such as *Lactobacillus* has been found to be a conserved feature (Salminen and Isolauri, 2006). However, we found *Bifidobacterium* in the cubs’ gut microbiota, which was rarely reported in previous studies (Tun et al., 2014, Peng et al., 2016). There are many reasons for the difficulty in detecting *Bifidobacterium*, such as the detection value being lower than the detection limit (strain-level genotyping technology would improve this) (Avershina et al., 2018). *Lactobacillus* and *Bifidobacterium* can inhibit the growth of pathogens by reducing the pH (by producing lactic and acetic acid) (Adams and Hall, 1988) or by competing for nutrients and epithelial adhesion sites (Agostoni et al., 2004). These bacteria play a vital role in cubs’ digestion of dairy products and absorption of nutrients. Our study also revealed that the juveniles’ dominant genera were *Streptococcus* and *Escherichia-Shigella*. *Escherichia-Shigella* can cause potentially fatal gastrointestinal tract disease in giant pandas (Zou et al., 2018, Wang et al., 2013), which is the most common disease in giant pandas, and it is particularly associated with mortality in young giant pandas (Aziz et al., 2013). Notably, it is necessary to focus on monitoring the dynamics of *Escherichia-Shigella* for ex-situ conservation of the giant panda population. We found high relative abundances of *Pseudomonas*, *Clostridium*, *Lactobacillus*, *Enterococcus*, and *Acinetobacter* in giant pandas in all age groups except the cubs. These bacteria are associated with cellulose, hemicellulose, and lignin digestion. Furthermore, *Pseudomonas* is related to the detoxification of cyanide compounds in bamboo. In conclusion, as giant pandas age, not only do endogenous changes cause changes in gut microbial composition, but dietary adaptations to developmental stages also play a role in shaping the gut microbiota.

In this study, the adults had highest nuber in the gut microbiota, and the lowest diversity. The cubs had lowest nuber in the gut microbiota, the highest diversity. The geriatrics and juveniles had similar alpha diversity. The results are consistent with findings in red pandas (Williams et al., 2018) and Shiba Inu dogs (Mizukami et al., 2019). A study on rhesus monkeys of different ages reported no difference in alpha diversity of the gut microbiota (Yatsunenko et al., 2012). However, it is generally believed that, in giant pandas, the alpha diversity of the gut microbiota in cubs is lower than that in adults (Janiak et al., 2021, Koenig et al., 2011). At different developmental stages, giant pandas have different feeding habits. During the cub stage, some cubs ate dairy products, while others also ate bamboo. Cubs’ microbiota is in a stage of colonization that is unstable and may be affected by exogenous factors, such as food resources. The adults ate bamboo almost exclusively. Adults’ gut microbiota tends to stabilize and can even be restored after perturbation. These factors may affect the diversity and abundance of the gut microbiota. Studies have suggested that, for species with different feeding habits at different developmental stages, differences in food resources are one of the most direct and important exogenous factors affecting the diversity and abundance of the gut microbiota (Wong et al., 2015, Vences et al., 2016, Zhang et al., 2018a, Zhao et al., 2018).Many studies have reported associations between age and gut microbiota diversity in giant pandas. Consistent with our findings, which showed that cubs, Guo et al. reported that the Shannon diversity index was significantly lower in the adults than in the cubs (Guo et al., 2020). Peng et al. reported that the alpha diversity of adults was significantly higher than that of geriatrics but not significantly different from juveniles (Peng et al., 2016). Another study reported that the gut microbiota diversity of adults was lower than that of geriatrics (Tun et al., 2014). A study exploring the relationships between age, season, and gut microbes in giant pandas showed that the alpha diversity of adults in autumn was similar to that of cubs (Xue et al., 2015). The reasons for the differences among studies regarding the age-related changes in the gut microbiota diversity of giant pandas may be as follows: (1) small sample size; (2) different habitats; (3) different sample processing (blending/vortexing); (4) different DNA extraction methods; (5) different amplification primers; and (6) different sequencing methods and platforms (Brooks et al., 2015, Wagner Mackenzie et al., 2015, Williams et al., 2016). In our study, the number of samples in the four age groups was relatively large and balanced. The sampling time was within a small window (April), the samples were distributed across Sichuan, the captive management methods were relatively consistent, and the bamboo species serving as staple food were relatively similar.

A comparison of beta diversity among the groups showed that the gut microbiota in the cubs was significantly different from that in the juveniles, adults, and geriatrics. The ANOSIM and Adonis results consistently showed that the between-group difference was greater than the within-group difference, indicating that our grouping strategy was appropriate. NMDS analysis divided the four groups of gut microbes into two parts: the cub and non-cub (juvenile, adult, and geriatric) gut microbiota, with a clear demarcation line between the two parts. The PLS-DA results support the NMDS conclusions. Therefore, we can divide the structure of giant pandas’ gut microbiota into two types: cub-type and non-cub-type (juvenile, adult, and geriatric) gut microbiota. From birth to 1.5 years old (cub stage), the diet involves dairy products and physiological function is limited, but after 1.5 years old (juvenile stage), more bamboo products are eaten and full physiological function begins to be developed. Therefore, it is inferred that the age-related changes in giant pandas represent a major factor in shaping the gut microbiota.

## 5 Conclusion

Our study indicates that the gut microbial community composition, abundance, and functional pathways differ among four age groups of giant pandas. In particular, the cubs and adults showed unusual alpha diversity (the adults had the highest richness, and the lowest diversity and homogeneity; the cubs had the lowest richness, the highest diversity, and the highest homogeneity, respectively), which may be caused by the cubs’ diet of dairy products with the addition of some bamboo. However, the specific causes need to be further verified by controlling for diet and age in future research. In addition, among the gut microbes of the cubs and juveniles, *Escherichia-Shigella* was a dominant genus. *Escherichia-Shigella* is a type of bacteria that causes gastrointestinal tract disease. It can even cause the death of young giant pandas. Therefore, during ex-situ population management of giant pandas, it is necessary to improve the monitoring of intestinal microorganisms in young giant pandas.

## 6 Acknowledgments

We thank Zong Cheng, Wei Ming, Yin Yuzhong, Yu Jianqiu, and others for their valuable advice and assistance with sampling. Special thanks go to China Conservation and Research Center for Giant Pandas (Dujiangyan, Sichuan Province, China), Chongqing Zoological Garden (Chongqing, China), and Chengdu Zoological Garden (Chengdu, Sichuan Provence, China) for their support.

## 9 Ethics Statement

Fecal samples were collected from China Conservation and Research Center for Giant Pandas Bifengxia Base (Ya’an, Sichuan Province, China), Dujiangyan Base (Dujiagnyan, Sichuan Province, China), Shenshuping Base (Wolong, Sichuan Province, China), and Chongqing Zoo (Chongqing, China). All sample collection procedures followed the guidance of the Sichuan Animal Society Ethics Committee and had no impact on animal health. All the samples were collected by an experienced laboratory technician and were immediately frozen in a liquid nitrogen container before being stored at −80°C. All animal work was approved in accordance with the EU Directive 2010/63/EU for animal experiments. All experiments were performed in line with the approved guidelines and regulations.

## 10 Author Contributions

HL contributed significantly to the conception of the work and drafted the manuscript. KL and GL were in charge of collecting samples. TL and HL participated in the sample collection and drawed the figures. CL participated in the sample collection and analysis of data for the work. QX, HL, and XZ participated in the sample collection. WW and YW analysed the data. GL, MG and GZ are the corresponding authors, conceived and participated in the design and coordination of the study, and helped draft the manuscript. Drafting the work or revising it critically for important intellectual content. or the acquisition, analysis or interpretation of data for the work. All authors read and approved the final manuscript.

